# Humanized CB1R and CB2R yeast biosensors enable facile screening of cannabinoid compounds

**DOI:** 10.1101/2022.10.12.511978

**Authors:** Colleen J. Mulvihill, Josh Lutgens, Jimmy D. Gollihar, Petra Bachanová, Edward M. Marcotte, Andrew D. Ellington, Elizabeth C. Gardner

**Affiliations:** Department of Molecular Biosciences, Center for Systems and Synthetic Biology, The University of Texas at Austin, Austin, TX 78712 USA; Antibody Discovery and Accelerated Protein Therapeutics, Center for Infectious Diseases, Houston Methodist Research Institute, Houston, TX 77030 USA; Genetic Design and Engineering Center, Department of Bioengineering, Rice University, 6100 Main St., Houston, TX 77005

**Keywords:** cannabinoid receptors, humanized yeast, G protein-coupled receptors

## Abstract

Yeast expression of human G Protein Coupled Receptors (GPCRs) can be used as a biosensor platform for the detection of pharmaceuticals. The Cannabinoid receptors type 1 and 2 (CB1/2R) are of particular interest, given the cornucopia of natural and synthetic cannabinoids being explored as therapeutics. We show for the first time that engineering the N-terminus of CB1R allows for efficient signal transduction in yeast, and that engineering the sterol composition of the yeast membrane optimizes CB2R performance. Using the dual cannabinoid biosensors, large libraries of synthetic cannabinoids and terpenes could be quickly screened to elucidate known and novel structure-activity relationships, including compounds and trends that more selectively target each of the two receptors. The biosensor strains offer a ready platform for evaluating the activity of new synthetic cannabinoids, monitoring drugs of abuse, and developing molecules that target the therapeutically important CB2R receptor while minimizing psychoactive effects.

## Introduction

GPCRs are common therapeutic targets, and an estimated 30% of FDA-approved drugs target this family of receptors^1^. The native pheromone response pathway of *Saccharomyces cerevisiae* is regulated by an endogenous GPCR (Ste2/Ste3)^2^, and the yeast receptors can be replaced with human GPCRs, leading to signal transduction via modified Gpa1 Gα proteins and activation of the MAPK cascade, ultimately inducing gene expression via the transcription factor Ste12.^3^

GPCRs of particular medical and industrial importance are the Cannabinoid Receptors, Type 1 (CB1R) and Type 2 (CB2R). The most abundant GPCR in the brain^4^, CB1R is activated by the psychoactive drug tetrahydrocannabinol (THC), but is also the target of endocannabinoids 2-arachadonoylglycerol (2-AG) and anandamide (AEA)^5^. These neurotransmitters exist as lipid precursors embedded in cell membranes where they are cleaved by lipases and liberated for receptor activation^6^. Endocannabinoid regulation *via* CB1R is implicated in neuronal excitability, where retrograde transmission of endocannabinoids from postsynaptic cells activates CB1R on presynaptic neurons and negatively regulates presynaptic neurotransmission via the Gα_i/o_ pathway^7^. CB1R dysregulation is, in turn, associated with schizophrenia^8^. In contrast to CB1R, the second of the two cannabinoid receptors, CB2R, is primarily expressed in leukocytes, where it regulates immune function. CB2R activation is broadly associated with an anti-inflammatory effect where CB2^−/−^ mice exhibit increased leukocyte recruitment and inflammatory marker production^9^. CB2R is not linked to psychoactivity and is a promising drug target for inflammatory diseases including arthritis, atherosclerosis, and inflammatory bowel disease^9^.

Synthetic cannabinoids have been developed to elicit a response from one or both cannabinoid receptors. For decades these drugs have been sold illicitly to consumers looking for similar psychoactive effects as THC. They are quite popular, being the second most-used illegal substance by young adults^10^. Unlike THC, these compounds are typically not identified in conventional drug screens. Many synthetic cannabinoids are much tighter binders to cannabinoid receptors than THC, and are often sold as mixtures uncharacterized for human use^10^. Together, the high potency and lack of regulation of these compounds have led to many cases of adverse effects from recreational use, including acute psychosis, seizures, dependence and death^10–13^. Governments have attempted to regulate these compounds, but regulation remains a challenge as new compounds and analogs of existing ones are created frequently that evade restriction^10^.

Given the role of cannabinoids in the treatment of chronic pain, epilepsy, and psychiatric disorders, there is a growing demand for next generation cannabinoid medicines. Ideal therapeutic candidates should activate CB2R while avoiding potent activation of CB1R and triggering subsequent psychoactive effects. Discrimination between the two receptors is challenging, as CB1R and CB2R share a high degree of sequence similarity, including almost identical binding pockets^14^.

Cannabinoid biosensor yeast strains have the potential to serve as a rapid, inexpensive, and robust screening platform. Unfortunately, they have not yet enabled facile comparisons between CB1R and CB2R activation. Herein, we demonstrate the engineering of CB1R and CB2R yeast biosensors by combining synthetic biology approaches that target receptor trafficking and membrane composition. Using these optimized strains, we screened more than 400 synthetic cannabinoids and terpenes, characterizing known effectors and discovering unknown functions of cannabinoids, including analogs of controlled drugs of abuse. The dual cannabinoid biosensors provide a rapid functional screen that can be used to readily rationalize structure-activity relationships at each receptor, and should accelerate the development of safe cannabinoid therapeutics into the future.

## Results and Discussion

### Engineering cannabinoid receptor function in yeast

While functional human GPCRs have previously been expressed in yeast, this had proved challenging for CB1R. To enable function, Ste2 (the native yeast GPCR of the *MAT*a haplotype), the negative regulator SST2 (a GTPase activating protein), and FAR1, which regulates cell cycle, were deleted in a manner consistent with prior efforts^15^. To efficiently transduce signals from the human machinery to the yeast pathway, the yeast 5 C-terminal residues of the G-alpha gene GPA1 were replaced with the sequence of the G_ai3_ human G-alpha variant as previously described^3^. We used ZsGreen1 to report on the activity of the pathway, integrating it into the genome in place of the pheromone-inducible Fig1 gene (**Fig. 1A**).

**Figure 1.**
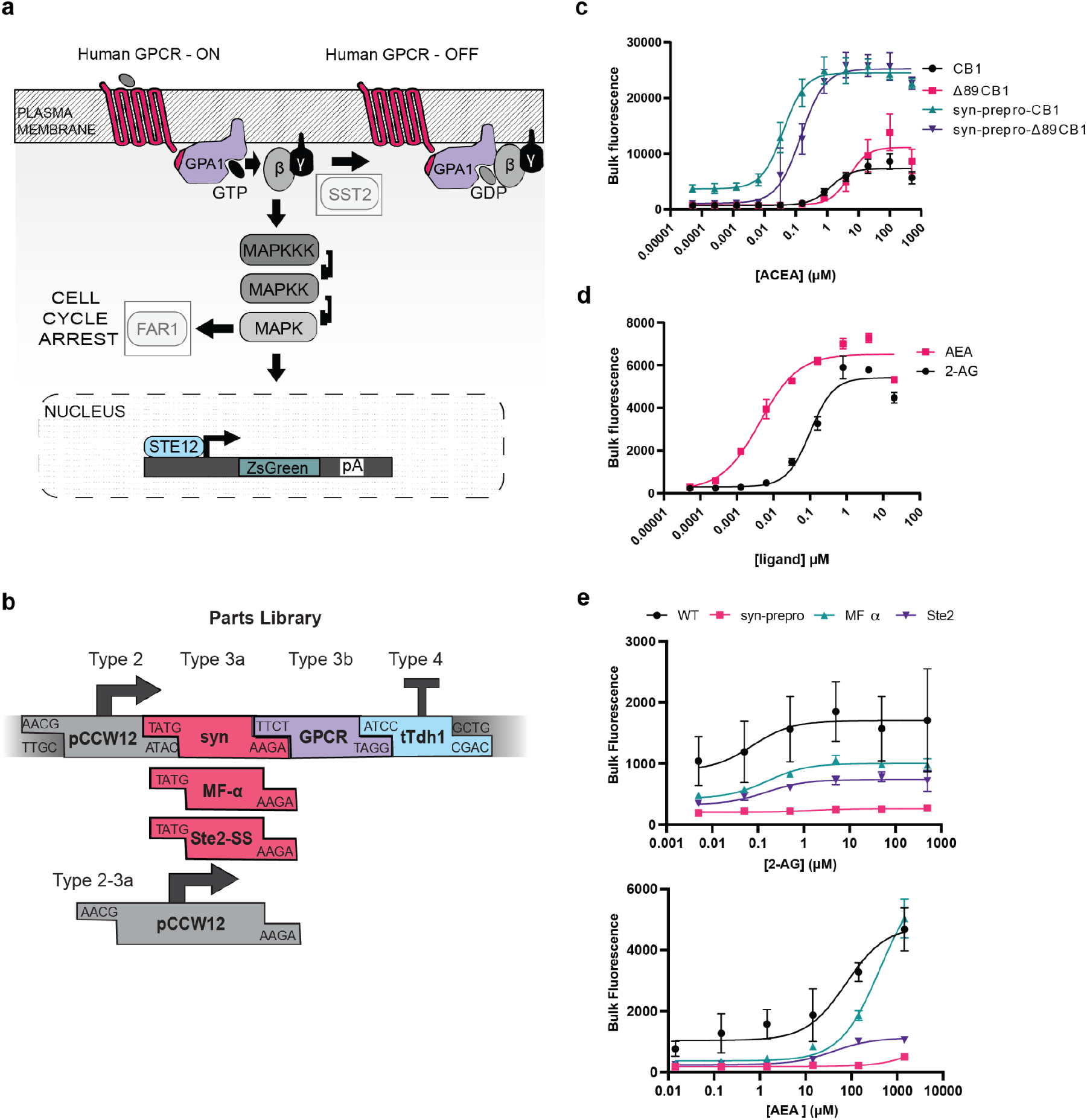
Development of CB1R and CB2R biosensors. (A). The yeast pheromone response pathway was genetically modified for human receptors. Greyed out boxes represent knocked out genes. The yeast pheromone GPCR Ste2 was replaced with a human GPCR. The Fig1 pheromone response gene was replaced with a ZsGreen reporter via knock-in. (B) The parts library for rapid, modular construction of trafficking-optimized GPCRs. Syn = synthetic pre-pro, MF-α = mating factor alpha pre-pro, Ste2-SS = Ste2 signal sequence. (C) ACEA dose response with CB1R. CB1, EC_50_=1.30 μM, 95% CI 0.80 to 2.12 μM. Δ89-CB1R, EC_50_ 5.05 μM, 95% CI 2.64 to 9.29 μM. syn-prepro CB1R EC_50_=0.04 μM, 95% CI 0.03 to 0.05 μM. Syn-prepro Δ89-CB1R, EC_50_=0.15 μM, 95% CI 0.10 to 0.21 μM. (D) Dose response of endocannabinoids 2-arachidonoyl glycerol (2-AG) and N-arachidonoylethanolamine (anandamide, AEA) with CB1R. AEA, EC_50_: 4 nM, 95% CI 2.62×10^−9^ to 6.92×10^−9^ M) than 2-AG (EC_50_: 100 nM, 95% CI 6.60×10^−8^ to 1.48×10^−7^M). (E) Dose responses with 2-AG and AEA, CB2R. 2-AG: WT, EC_50_=0.07 μM, 95% CI 0.0006881 to 3.70 μM; syn-prepro, EC_50_=1.54 μM, 95% CI 0.02 to 70.9 μM; MF alpha, EC_50_=0.16 μM, 95% CI 0.08 to 0.32 μM; Ste2, 0.15 μM, 95% CI 0.39 to 0.58 μM. AEA: WT, EC_50_=76.9 μM, 95% CI 18.6 to 242 μM; syn-prepro, EC_50_=7620 μM, 95% CI 630 to +infinity μM; MF alpha, EC_50_=419 μM, 95% CI 278 to 693 μM; Ste2, 40.8 μM, 95% CI 29.3 to 56.8 μM. For all experiments, *n*=3. Nonlinear regression was fit as a 4-parameter logistic equation.

Initially, the wild-type, leaderless CB1R showed low (<4-fold) signaling with the synthetic cannabinoid ACEA (**Fig. 1C**), which we speculated might be due to defects in plasma membrane localization. To improve basal function, we modified the popular MoClo yeast toolkit parts library^16^ to include yeast-optimized pre-pro signals (designated Type 3a) after the Type 2 promoter and before the Type 3 GPCR, yielding seamless fusions (**Fig. 1B**). Human CB1R also has a long N-terminal domain that can block co-translational insertion into the endoplasmic reticulum^17^, thus partially re-directing the receptor to localize in the mitochondria^18^. To better specify transport, we created a N-terminally truncated version of the CB1 receptor, both with and without the syn-prepro secretion sequence. The syn-prepro modification alone improved activation to approximately 7-fold, while the truncated receptor showed 15-fold activation; both modifications together yielded 28-fold activation. However, this improved dynamic range came at a cost: the syn-prepro-Δ89-CB1R was ultimately found to have an EC_50_ of 147 nM, a 3.8-fold drop in sensitivity compared to the full-length, syn-prepro construct (**Fig. 1C**). This tradeoff may be due to an alteration in conformational equilibria in the deletion variant, since syn-prepro-Δ89-CB1R also showed a higher fraction of signaling cells in the population (**Supp. Fig. 1**). The CB1R biosensor strain was also benchmarked with the endocannabinoids 2-AG and AEA (**Fig. 1D**), and was found to have sensitivities (AEA EC_50_ of 4 nM; 2-AG EC_50_ of 100 nM) in accord with previously published values^19^. The dramatic functional improvement achieved by the combination of N-terminal truncation and addition of a synthetic pre-pro leader highlights that proper receptor trafficking can be an issue when expressing GPCRs in yeast heterologously. Since dynamic range was likely to be of greatest utility for screens, especially with compounds with otherwise unknown affinities, we chose the combination of syn-prepro-Δ89-CB1R with ACEA to serve as the benchmark for further studies.

Unlike CB1R, CB2R is functional in yeast in its native form^20,21^. To further optimize function, we compared three leader peptides to the native N-terminal sequence (**Fig. 2**), assaying the sensor’s performance with the endocannabinoids AEA and 2-AG. The MFa leader showed the greatest dynamic range in signaling with AEA (22-fold ± SD 3.2, WT 9.6 ± SD 0.8). The improvement in dynamic range again generated a tradeoff in performance, as the wild-type receptor was again more sensitive with both agonists than the engineered variant. Signaling of all receptor versions showed greater sensitivity with 2-AG relative to AEA, concordant with previous findings in mammalian cells that CB2R is more sensitive to 2-AG than AEA^22^. Again, since dynamic range was of greatest import for maximizing screening capabilities, we selected the MFa-CB2R construct and AEA moving forward.

**Figure 2.**
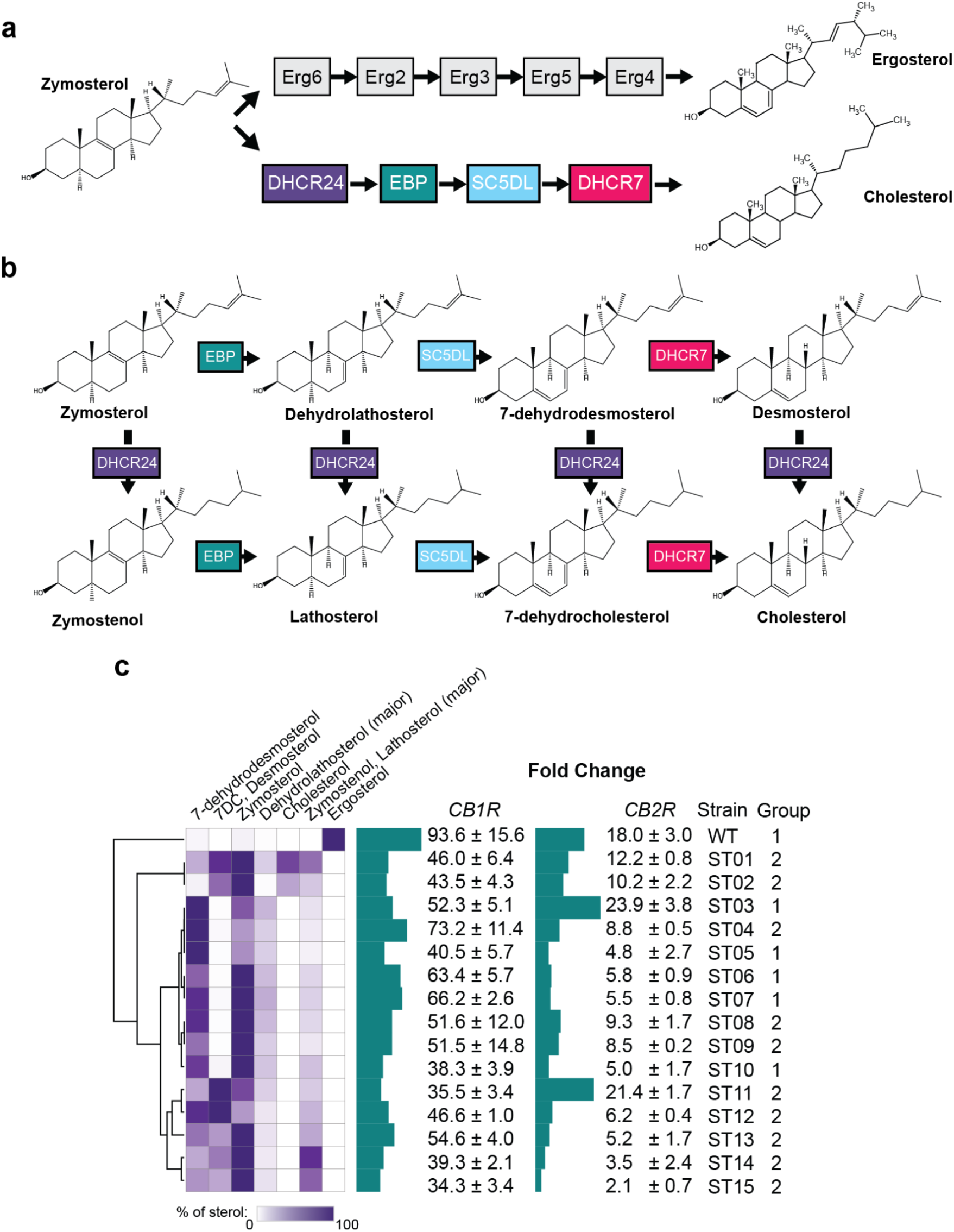
Optimization of sterol environments for CB1R and CB2R biosensors. (A) Comparison of ergosterol and cholesterol production pathways in yeast and humans, respectively, after the common intermediate zymosterol. (B) Cholesterol biosynthetic intermediates and the enzymes that act on them after zymosterol. (C) Fold change in signaling results for CB1R and CB2R in sterol-modified strains. Strains were engineered to produce cholesterol and\or cholesterol biosynthetic intermediates, and sterol content was measured using GC-MS in a previously published work reproduced here with permission. Dose responses were performed for both receptors in each strain, and fold changes of maximum signal over signal with 0 μM agonist were calculated. The agonist ACEA was used for CB1R, and AEA was used for CB2R. For all experiments, *n*=3. Nonlinear regression was fit as a 4-parameter logistic equation.

In addition, despite the fact that human GPCRs typically work in yeast at neutral pH^23–25^, CB2R showed prohibitively strong constitutive signaling at pH 7.1 (a phenomenon also previously noted with the heterologously expressed serotonin receptor^26^). When the biosensor was assayed at pH 5.8, we observed greatly reduced background with both AEA and 2-AG. Therefore, we subsequently screened all compounds with the CB2R biosensor strain at pH 5.8.

### Refactoring sterol composition can enhance cannabinoid receptor activity

Cholesterol is necessary for the function of many GPCRs^27^ (with ~40% of GPCR PDB structures being co-crystallized with cholesterol^28^), and providing cholesterol or intermediate metabolites to yeast can dramatically impact receptor function^24^. Since CB1R is known to contain cholesterol recognition amino acid consensus (CRAC) motifs^29^, we hypothesized that introducing the cholesterol biosynthetic machinery into CB1R- and CB2R-expressing biosensor strains might further improve receptor function.

Yeast and human sterol biosynthesis share a common zymosterol precursor that can be converted to ergosterol through a five enzyme pathway or to cholesterol through a four enzyme pathway (**Fig. 2A**). The four-enzyme pathway produces seven metabolites (including cholesterol) in addition to zymosterol (**Fig. 2B**). We previously created multiple yeast strains that produced varying levels of these seven metabolites^24^.

The syn-prepro-Δ89-CB1R expression construct was transformed into fifteen strains selected to cover the metabolic space. Previously, cholesterol was shown to negatively regulate CB1^29^ receptors, although it was unclear if ergosterol would have the same effect, as it differs by two double bonds and a methyl group. Dose-responses with ACEA yielded a striking pattern in which all cholesterol intermediates were found to have a deleterious effect on CB1R signaling compared to ergosterol only (**Fig. 2C**), with strains containing cholesterol or any of its intermediates exhibiting lower sensitivities and dynamic ranges in signaling (**Supp. Fig. 2**). We thus selected the wild type, ergosterol-producing strain for subsequent studies, as it showed both the highest fold-change and most sensitive signaling (EC_50_ of 51 nM, **Supp. Fig. 2**).

Similarly, we transformed MFa-CB2R into the fifteen strains with distinct sterol environments and determined dose-responses using AEA (**Fig. 2C**). Unlike CB1R, the literature is mixed regarding cholesterol’s effect on CB2R function: Prior studies have shown that depletion of cholesterol has no effect on 2-AG binding to CB2R, but also that cholesterol increases constitutive activity of the receptor and alters ligand binding profiles^30,31^.

Some of the strains we tested (ST01, ST02, ST04, ST09, ST11, ST12, and ST13) showed low responsivity until maximal concentrations of AEA were added, making EC50 determinations impossible (**Supp. Fig. 3**). Other strains (ST03, ST05, ST06, ST07, and ST10) proved more sensitive to AEA than the wild-type with ergosterol, but showed a dip in signaling at the highest AEA concentrations (**Supp. Fig. 3).** Interestingly, this latter phenomenon has been observed previously with other CB2R agonists in yeast^20^, and is consistent with known functional interactions between cholesterol and cannabinoids. We suspect that compound interactions at high concentrations with cholesterol and cholesterol-like membrane components may alter membrane fluidity or other features in a way that is deleterious to receptor function^32^.

For CB2R, we did not observe a clear distinction between the membrane compositions of the two classes of strains, nor was it obvious which specific set of membrane components led to improved signaling, although there was a general trend between reduced signaling and the presence of the cholesterol-adjacent compounds 7-dehydrocholesterol and desmosterol. This lack of clarity emphasizes the need for a screening-based approach to membrane engineering, as we have previously shown with other human GPCRs^24^. For subsequent experiments, we opted to use the ST03 strain as it showed the highest dynamic range and was also sensitive at lower concentrations of AEA, with a greater signal at lower concentrations of AEA compared to background than the wild type ergosterol-producing strain (**Supp. Fig. 3**).

### High-throughput drug screening of cannabinoids with dual cannabinoid receptor biosensor strains

The availability of two biosensor strains with optimized cannabinoid receptor function immediately presents an opportunity to determine the relative activities of a variety of cannabinoids and other compounds. A compound library of over 300 synthetic cannabinoids was screened against both strains; as a negative control, DMSO or the inverse agonist rimonabant was included on each plate analyzed by flow cytometry. The compound library included known characterized agonists and or structurally similar compounds; for example, AB-PINACA, ADB-PINACA, and AB-FUBINACA are all structurally related Schedule I controlled substances found in synthetic cannabis products. All test compounds were delivered at 1 μM, orders of magnitude in excess of typical EC_50_ values for agonists, in order to identify even low affinity interactions. To better identify highly active compounds, assays were also carried out at 10 nM concentrations.

The biosensor screening assay showed that a large proportion of the compounds were active with the CB1R receptor relative to ACEA (**Fig. 3A**), and many compounds showed higher activity with CB2R than an endocannabinoid control, AEA (**Fig. 3B)**. To validate the results, known and predicted agonists and antagonists of the receptors were examined individually. The known antagonist URB447^33^, predicted antagonist WIN54,461^34^, non-binder HU-211^35^, cannabidiol degradation product HU-311^36^, or selective CB2R ligands A-796260^37^, AM-1241^38^, MDA 77^39^ all showed less than 25% relative fluorescence of the ACEA positive control at CB1R at 10 nM (**Supplemental Figure 4 A**), indicating that the yeast biosensors could readily identify both agonists and antagonists. In contrast, known agonists AB-FUBINACA, ADB-PINACA, and AB-PINACA showed 79-84% maximal activity at this concentration.

**Figure 3.**
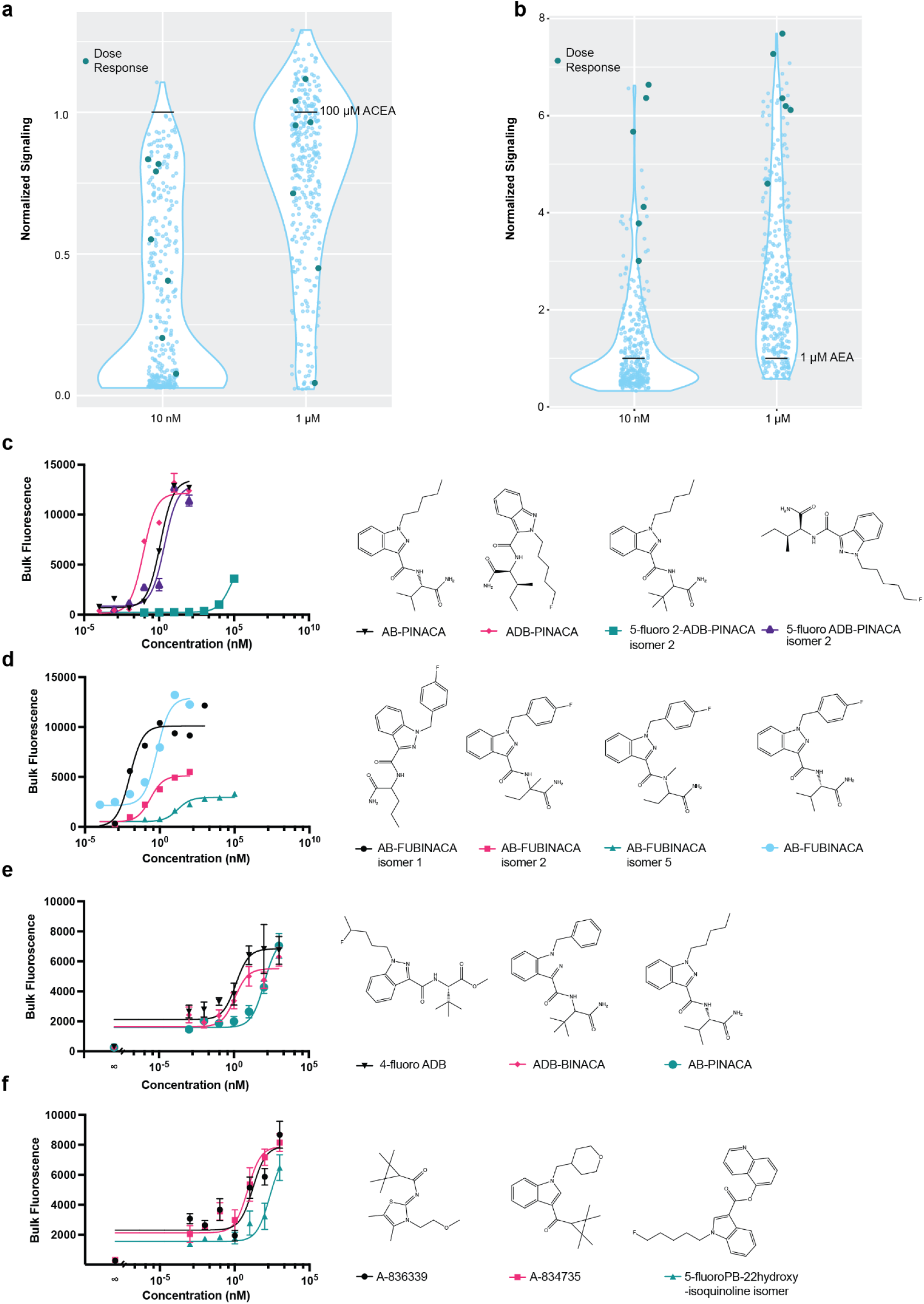
Screen of cannabinoid receptors with a synthetic cannabinoid compound library. (A) Screen of CB1R with synthetic cannabinoid compounds at 1 μM and 10nM concentrations. Signaling was normalized to signaling with 100 μM ACEA, set at “1”. “Dose response” compounds were those chosen for further analysis by dose response. (B) Screen of CB2R with synthetic cannabinoid compounds at 1 μM and 10nM concentrations. Signaling was normalized to signaling with 1 μM AEA, set at “1”. “Dose response” compounds were those chosen for further analysis by dose response. (C) Dose responses with FUBINACA compounds. AB-FUBINACA, EC_50_=0.66 nM, 95% CI 0.35 to 1.14 nM; AB-FUBINACA isomer 1, EC_50_=0.01 nM, 95% CI 0.01 to 0.02 nM; AB-FUBINACA isomer 2, EC_50_=0.23 nM, 95% CI 0.1356 to 0.42 nM; AB-FUBINACA isomer 5, EC_50_=15.86 nM, 95% CI 7.63 to 38.36 nM. (D) Dose responses with PINACA compounds. AB-PINACA, EC_50_=1.21 nM, 95% CI 0.96 to 1.52 nM; ADB-PINACA, EC_50_=0.09 nM, 95% CI 0.05 to 0.14 nM; 5-fluoro ADB-PINACA isomer 2, EC_50_=2.25 nM, 95% CI 1.26 to 3.97 nM. (E) Dose responses with select synthetic cannabinoids. A-836339, EC_50_=16.3 nM, 95% CI 3.53 to 186 nM; A-834735, EC_50_=7.42 nM, 95% CI 2.12 to 22.2 nM; 5-fluoro PB-22 5-hydroxyisoquinoline isomer, EC_50_=216 nM, 95% CI 66.3 to 1300 nM. (F) Dose responses with select synthetic cannabinoids. 4-fluoro ADB, EC_50_=1.35 nM, 95% CI 0.274 to 5.03 nM; ADB-BINACA, EC_50_=1.15 nM, 95% CI 0.254 to 4.87 nM; AB-PINACA, EC_50_=107 nM, 95% CI 42.1 to 251 nM.

The CB1R biosensor gave consistent characterizations within compound classes. For example, in the PINACA agonist series (AB-PINACA, ADB-PINACA AND 5-fluoro ADB-PINACA isomer 2) all were shown to have EC_50_ values in the low nanomolar range (**Figure 3C**). Dose-response curves for AB-FUBINACA isomers were also obtained, including for many compounds that had previously been uncharacterized. AB-FUBINACA and AB-FUBINACA isomer 1 showed high maximal signaling, while isomers 2 and 5 had a lower response (**Figure 3D**). Since AB-FUBINACA has been implicated in overdoses from recreational use along with a number of its variants^11^. knowledge of the activity of structural variants could be useful for determination of compound scheduling.

For CB2R, there was again consistency within compound classes, and there were frequently structural cognates for activity. For example, structural analogues, such ADB-PINACA and AB-PINACA, showed high activity (**Supp. Table 1)**, with the phenylmethyl group at the 1H-indazole and dimethylpropyl of ADB-BINACA being preferred by roughly 100-times over the pentyl and isopropyl, respectively, of AB-PINACA (**Fig. 3E**). To our knowledge, ADB-BINACA had previously been uncharacterized against CB2R. The compound 4-fluoro ADB showed the greatest sensitivity amongst compounds tested (EC_50_ of 1.35 nM, **Fig. 3E**), is known to be highly active at cannabinoid receptors, and is implicated in at least one death from overdose^13^.

Similarly, the compounds A-836339 and A-834375 both have a tetramethylcyclopropylmethanone group adjacent to the carbonyl, which is known to confer CB2R bias and strong signaling^40^, while A-834735 and 5-fluoro PB22 share the indole moieties common to many synthetic cannabinoids. A-836339 and A-834375 had EC_50_s of 16.3 and 7.42 nM, respectively (**Fig. 3F**). Interestingly, the 5-fluoro PB-22 5-hydroxyquinoline isomer is an otherwise uncharacterized isomer of 5-fluoro PB-22 and showed robust signaling with an EC_50_ of 216 nM (**Fig. 5C**).

### High-throughput drug screening of terpenes with dual cannabinoid receptor biosensor strains

It has been hypothesized that THC and many of the terpenes found in cannabis strains work synergistically to create strain-variant effects^41^, so we also assessed a library of terpenes against both the CB1R and CB2R biosensor strains (**Fig. 4A, B**). A number of compounds in the library have been found in cannabis, and others are known to be bioactive^41^. In order to detect even minor activities, compounds were screened at 100μM, but very few compounds showed activity against either receptor. Amongst terpenes found to activate CB1R, both (+)-β-Citronellol and β-Eudesmol have previously been isolated from cannabis^42,43^, and β-Eudesmol showed signaling at lower (10μM) concentrations (**Fig. 4C**). Surprisingly, totatrol, isolated from the native New Zealand *Podocarpus totara* and more recently found in a wider range of plants^44^, was found to be active against CB2R at 10 nM, with an EC_50_ of 485 nM (**Fig. 4D**). While totarol has known therapeutic applications and antimicrobial properties^45,46^, we believe this to be the first documented activity against CB2R.

**Figure 4.**
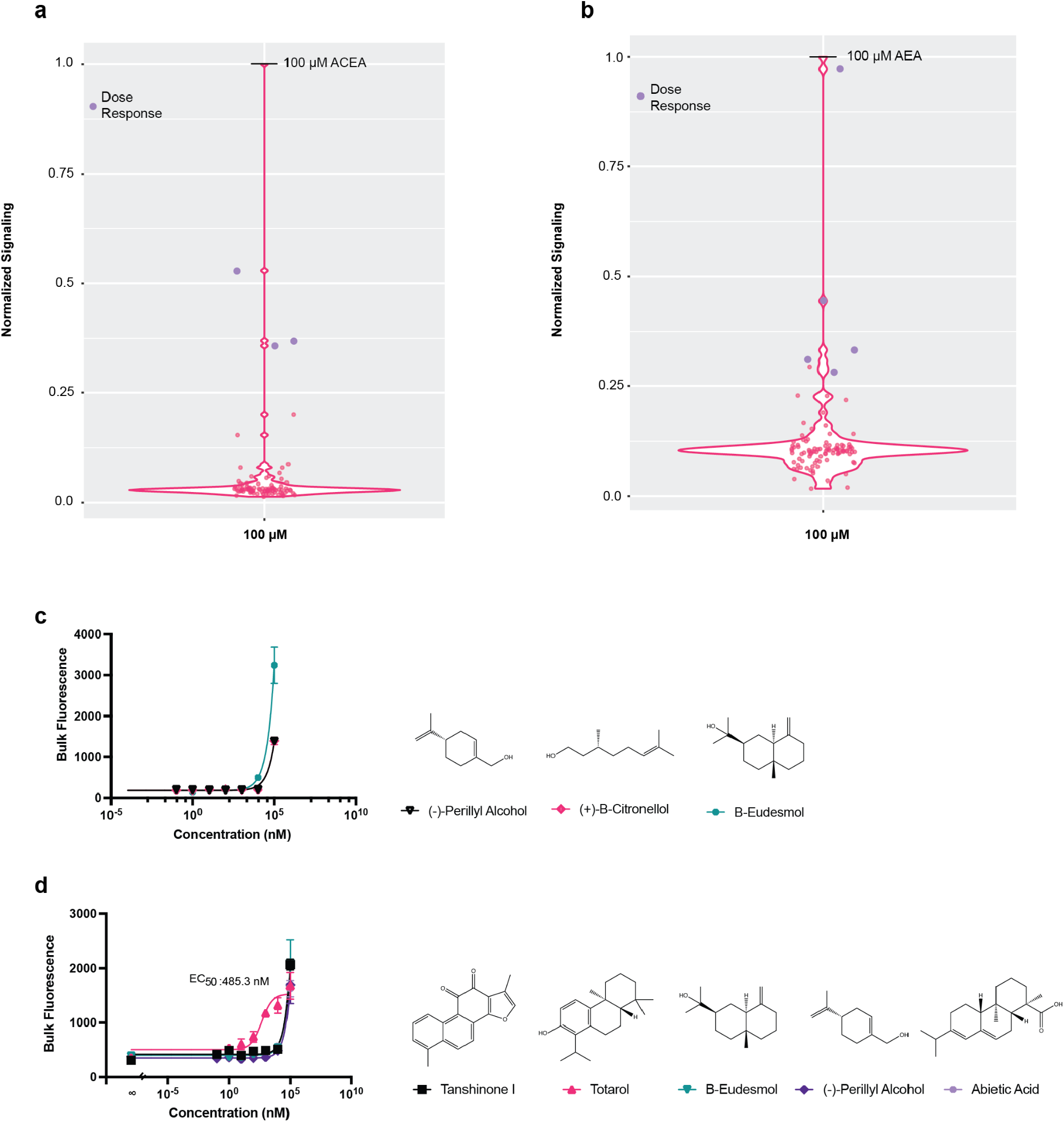
Screen of cannabinoid receptors with a terpene library. (A) Signaling at CB1R with terpene compounds at 100 μM. Signaling was normalized to signaling with 100 μM ACEA, set at “1”. “Dose response” compounds were those chosen for further analysis by dose response. (B) Signaling at CB2R with terpene compounds at 100 μM. Signaling was normalized to signaling with 100 μM AEA, set at “1”. “Dose response” compounds were those chosen for further analysis by dose response. (C) Dose responses at CB1R with compounds that demonstrated signaling in the terpene screen over 0.25 that of the 100 μM ACEA control. (D) Dose responses at CB2R with compounds that demonstrated signaling in the terpene screen over 0.25 that of the 100 μM AEA control with the exception of atractylenolide III. Totarol, EC_50_=485.3 nM, 95% CI 149.4 to 1443 nM. For all dose response experiments, *n*=3. Nonlinear regression was fit as a 4-parameter logistic equation. For the screens, *n*=1.

### Results with dual cannabinoid biosensor strains recapitulate and extend opportunities for receptor-specific targeting

Because the binding pockets of CB1R and CB2R are so similar, it has historically proven challenging to find compounds that activate one receptor but not both^14^. For example, the potent agonist 4-fluoro ADB shows high activity against both CB1R and CB2R^47,48^. For the compounds tested with the dual cannabinoid biosensor strains, the overall overlap in binding and activation between receptors is also apparent (**Fig. 5A**, which synthesizes the CB1R and CB2R results with 1μM compound from **Figs. 3A, B**; analyses with 10nM are in **Supp. Fig. 4 B)**. The ability of many compounds to activate or modulate CB1R recapitulates literature results^14^, and is consistent with the fact that CB1R is able to accomodate a wider variety of ligands than CB2R, due to the greater flexibility of its binding site across receptor activity states^14^.

**Figure 5.**
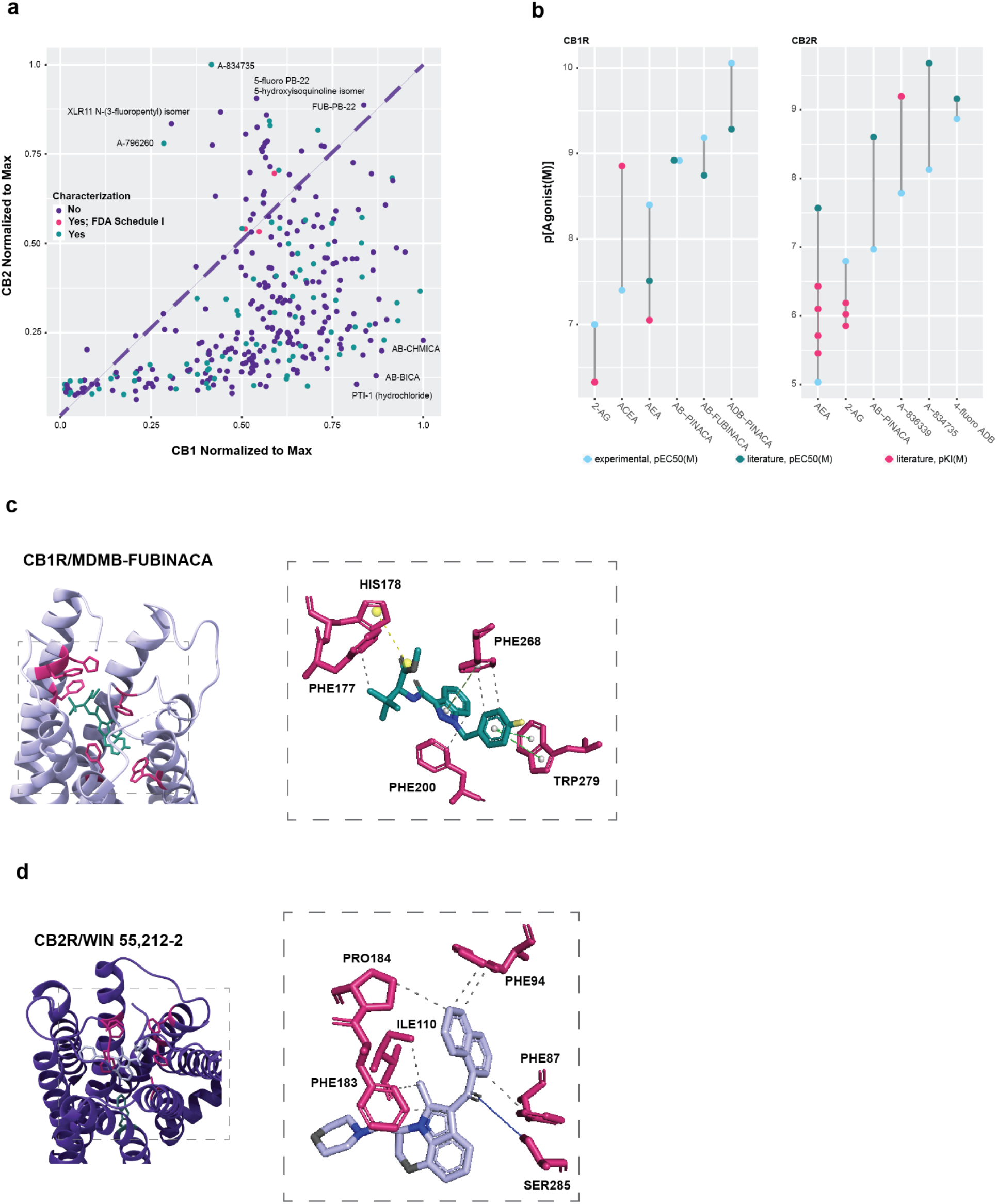
Results with dual cannabinoid biosensor strains recapitulate and extend opportunities for receptor-specific targeting. (A) Synthetic cannabinoid compounds screened against CB1R and CB2R at 1 μM, normalized to the maximum signal for each receptor at this concentration. Large distances from the right and left of the plotted line connote rough bias towards CB1R and CB2R, respectively. Compounds are labeled based on if they, to our knowledge, have been characterized biochemically against one or both cannabinoid receptors. (B) Comparison of yeast experimental EC_50_ values with EC_50_ and Ki values found in the literature in mammalian cells^9,48,49,58–62^. When there were multiple experimental values for an agonist and receptor, the optimized sterol and leader background was chosen. (C) Rendering of previously published crystal structure CB1R PDB 6N4B^50^ bound to the agonist MDMB-FUBINACA. Residues expected to interact with the agonist are highlighted. Types of interactions are labeled. (D) CB2R PDB 6PT0^52^ bound to the agonist WIN-55,212-2. Residues expected to interact with the agonist are highlighted. Types of interactions are labeled. For (C) and (D), interactions were predicted using a previously described tool^63^. Nitrogens are highlighted on agonists in blue, oxygens in gray, and fluorine in yellow.

While the dual cannabinoid biosensor strains were largely consistent with the known literature, EC_50_ and K_i_ values were often higher than found in previous studies with mammalian cells (**Fig. 5B**). At least some of the differences observed may again be due to our observation that membrane composition can greatly impact receptor function: other groups have found evidence that cholesterol may act as a direct binding partner of endocannabinoids in the plasma membrane, and possibly have a role in membrane insertion of these compounds^32^, and thus the optimization of cholesterol levels may prove to be both receptor- and compound-specific. Nonetheless, there was a general concordance between the EC_50_ values for AB-PINACA with CB1R in yeast and in mouse AtT-20 neuroblastoma cells^49^ (both 1.2 nM), and for 4-fluoro ADB with CB2R in yeast (1.35 nM) and in (HEK) 293 T cells (0.69 nM^48^), and overall general trends for modulation appeared the same between yeast-based and mammalian assays.

The ability to rapidly carry out assays with directly comparable results, coupled with structural analyses, now allows us to better identify receptor-specific compounds. The previously determined crystal structure of CB1R co-crystallized with the synthetic cannabinoid MDMB-FUBINACA^50^ indicates that the diazole ring (a shared feature between MDMB-FUBINACA and the AB-PINACA analogs) creates hydrophobic contacts with F200 and F268, and positions additional hydrophobic contacts with F174, F177, H178, and F200 (**Fig. 5C**) that lead to rearrangement of TM2 of CB1R during receptor activation. In consequence, the relative affinities of ADB-PINACA isomers (**Fig. 3C; Supp. Table 1)** and AB-FUBINACA isomers (**Figure 3D)** can be rationalized, with the alkyl group adjacent to the amide bond differentially activating the receptor in several formats, including tert-butyl, 2-butyl, and isopropyl.

The high-throughput yeast-based assays also potentiate new forays into the identification of receptor-specific ligands. Amongst the CB1R agonists that were poor CB2R signalers (AB-BICA, AB-CHMICA and PTI-1), PTI-1 has a N1-indole pentyl side chain known to favor CB1R agonism^40^, similar to AB-PINACA and ADB-PINACA (**Figure 3C**)^51^. Interestingly, the uncharacterized fluoropentyl isomers of AB-PINACA roughly halved signaling with CB2R (**Supp. Table 1**), and further modulation of these moieties may lead to greater receptor discrimination. Discrimination *via* the amide-bonded moieties on synthetic cannabinoids (as described above) can also be rationalized by examining both structures and activities in the dual biosensor assay: AB-BICA is structurally similarly to ADB-BINACA, a strong CB2R agonist, but has a methylpropyl group adjacent to the amide rather than the dimethylpropyl of ADB-BINACA (**Supp. Fig. 5**), potentially indicating that CB1R favors the smaller carbon side group in this position.

Conversely, the drug A-796,260 was among the top CB2R targets that demonstrated low signaling with CB1R. This compound is a known CB2R-biased agonist whose 3-tetramethylcyclopropylmethanone group (in lieu of a 3-benzoyl or 3-naphthoyl group) confers CB2R bias^40^. Both XLR11 N-(3-fluoropentyl) isomer and A-834735 are also CB2R-biased, and, like A-796,260, contain a 3-tetramethylcyclopropylmethanone group. This group proved important for CB2R signaling but precluded full CB1R activity– eight of the top twenty compounds at CB2R had this structural group, but none of the top twenty compounds at CB1R did (**Figure 5A)**. A previously determined crystal structure of CB2R co-crystallized with the synthetic cannabinoid WIN 55,212-2 highlights that the bulky 3-indole adducts on these various compounds trigger conformational changes in CB2R via the toggle switch W258^52^ (**Fig. 5D**). In a similar vein, some thirteen hydroxyquinoline isomers of PB-22 were screened with variable results across the receptors. Nitrogens at the 8th and 5th positions led to the greatest activity with CB2R, and dramatic decreases in activity were observed at the 3rd, 6th, and 7th positions (**Supp. Table 2**), further indicating that this may be a valuable new class of compounds where reactivities with CB2R could be finely tuned.

## Conclusions

The cannabinoid receptors CB1R and CB2R are of particular clinical interest for their roles in pain relief, appetite stimulation or suppression, and anti-epileptic properties. By engineering CB1R and the membrane environment of CB2R, we were able to generate dual biosensors that could be used for the rapid screening of a wide variety of cannabinoids and other compounds, such as terpenes, including dozens of previously uncharacterized compounds. The results obtained with the yeast-based biosensors were congruent with those previously seen in mammalian cells, and further allowed the ready identification of known and new compounds with specificities for individual receptors. Based on functional and structural analyses we have now identified PTI-1 as a potentially useful CB1R agonist, and a series of PB-22 isomers for development as CB2R agonists. Small structural modifications were found to greatly impact relative receptor specificity, with some PINACA modifications greatly reducing activity with CB2R while leaving CB1R activity unafffected (**Supp. Table 1**). Similarly, some PB-22 hydroxyquinoline isomers favored CB1 by almost a factor of 3 (5-fluoro PB-22 4-hydroxyquinoline isomer; **Supp. Table 2**), while other closely related compounds favored CB2 by roughly 2-fold (5-fluoro PB-22 5-hydroxyisoquinoline isomer; **Supp. Table 2**).

In addition to advancing pharmaceutical drug development, the emerging cannabis industry exposes consumers to a large number of uncharacterized compounds. The sheer speed at which many psychoactive mixtures can be created makes it difficult for regulatory bodies and health authorities to keep pace, highlighting the need for studies that can quickly provide insights into receptor binding and specificity. The availability of facile comparative assays *via* the dual yeast biosensors builds on earlier work expressing CB2R in yeast^20,21^ and may potentially provide a straightforward basis for comparison suitable for both research and regulatory organizations.

Finally, given that the complete biosynthesis of cannabinoids has been achieved in yeast^53^, and that yeast biosensor strains expressing GPCRs have begun to show promise in detecting metabolites during bio-manufacturing^54,55^, we can readily envision adapting the GPCR-based sensors herein to the conjoined screening and selection of new pathways and receptors that are specific for any of a variety of cannabinoid and other compounds.

## Methods and materials

### Molecular biology

All cloning was performed using Golden Gate assembly following the MoClo Yeast toolkit^16^ with some adaptations (see Parts list). Assemblies were performed as follows: 20 fmol part plasmids, 10,000 units Type IIs restriction enzymes (T7 DNA ligase, Esp31/BsaI-v2, NEB), and 1 μL T4 DNA ligase (NEB) in a 10 μL reaction. Thermal cycling was performed as follows: 1 minute 37°C, 2 minutes 16°C, for 25 cycles, then 37°C for 30 minutes, and 80°C for 10 minutes. 5 μL each reaction was transformed into 100 μL DH10B and transformed according to the Mix and Go Transformation kit (Zymo Research).

### Yeast transformations

Yeast background strains were BY4741 (See yeast strain list). Yeast transformations were performed according to the EZ transformation II kit (Zymo research). 100 μL of cell prep was transformed with 1 μg or 5 μL of plasmid.

### Functional assays

Yeast colonies were picked and grown overnight to saturation in pH 5.8 SD-His media in a 2.2 mL deep well plate (Axygen) grown in a plate-shaking incubator (30°C, 1000 rpm, 3 mm orbital). In the morning cultures were diluted 1:10 or 1:25 in SD-His media buffered to pH 7.1 with 100 mM MOPSO (unless otherwise specified). Ligands were added to the culture and incubated with shaking for 8 hours. Cells were then washed three times with ice-cold Tris buffer (pH 7) and diluted 1:20 for cytometry. All cytometry was performed on a Sony SA3800 spectral analyzer. Each sample was tested for 10,000 events, and read at a rate of 1,000 events per second. All assays as reported here were filtered only for singlet populations.

## Acknowledgements

The authors thank Dr. Kevin Drew for assistance with molecular modeling and Dr. Katy Kao for technical assistance. Financial support was provided by a Cooperative Agreement between the University of Texas at Austin and DEVCOM Army Research Laboratory to A.D.E. and E.M.M. (W911NF-17-2-0091). E.M.M. acknowledges additional support from the Welch Foundation (F-1515), Army Research Office (W911NF-12-1-0390), and NIH (R35 GM122480). A.D.E. acknowledges support from the National Institute of Health (1R21AT010777-01) and the Welch Foundation (F-1654).

The authors understand that there are potential security implications to their work, and have consulted relevant security and law enforcement contacts about potential misuse of this research. The compound library available from Cayman Chemical does not require a DEA license^56,57^; regardless, this research was carried out in accord with DEA license RE0496895 (Schedule 1 compounds) and RE0562581 (Schedule 2 compounds).

## Author Contributions

Conceptualization: E.C.G., C.J.M. J.D.G.; Methodology: E.C.G., C.J.M., J.D.G.; Investigation: E.C.G., C.J.M, J.L.; Data Analysis & Interpretation: E.C.G, C.J.M., P.B., E.M.M., A.D.E.; Data curation: E.C.G, C.J.M., A.D.E.; Original draft: E.C.G, C.J.M.; Review & Editing: A.D.E, J.D.G, E.M.M.; Funding: A.D.E., E.M.M.

