## Supplementary Figure and Tables for "Humanized CB1R and CB2R yeast biosensors enable facile screening of cannabinoid compounds"

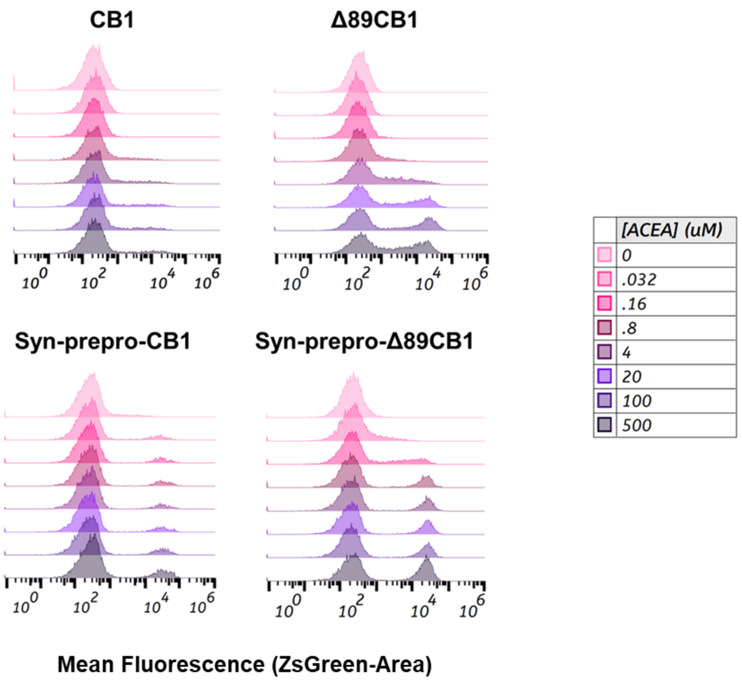


**Supplementary Figure 1. Fluorescence population shifts with CB1R variants.** Dose response of ACEA with four versions of the CB1 receptor: with and without the 89-residue amino terminal truncation, and with and without the synthetic pre-pro signal. Fluorescence was measured via cytometry, in which a population of singlets were binned for fluorescent response and total population ZsGreen signal (area) is shown.

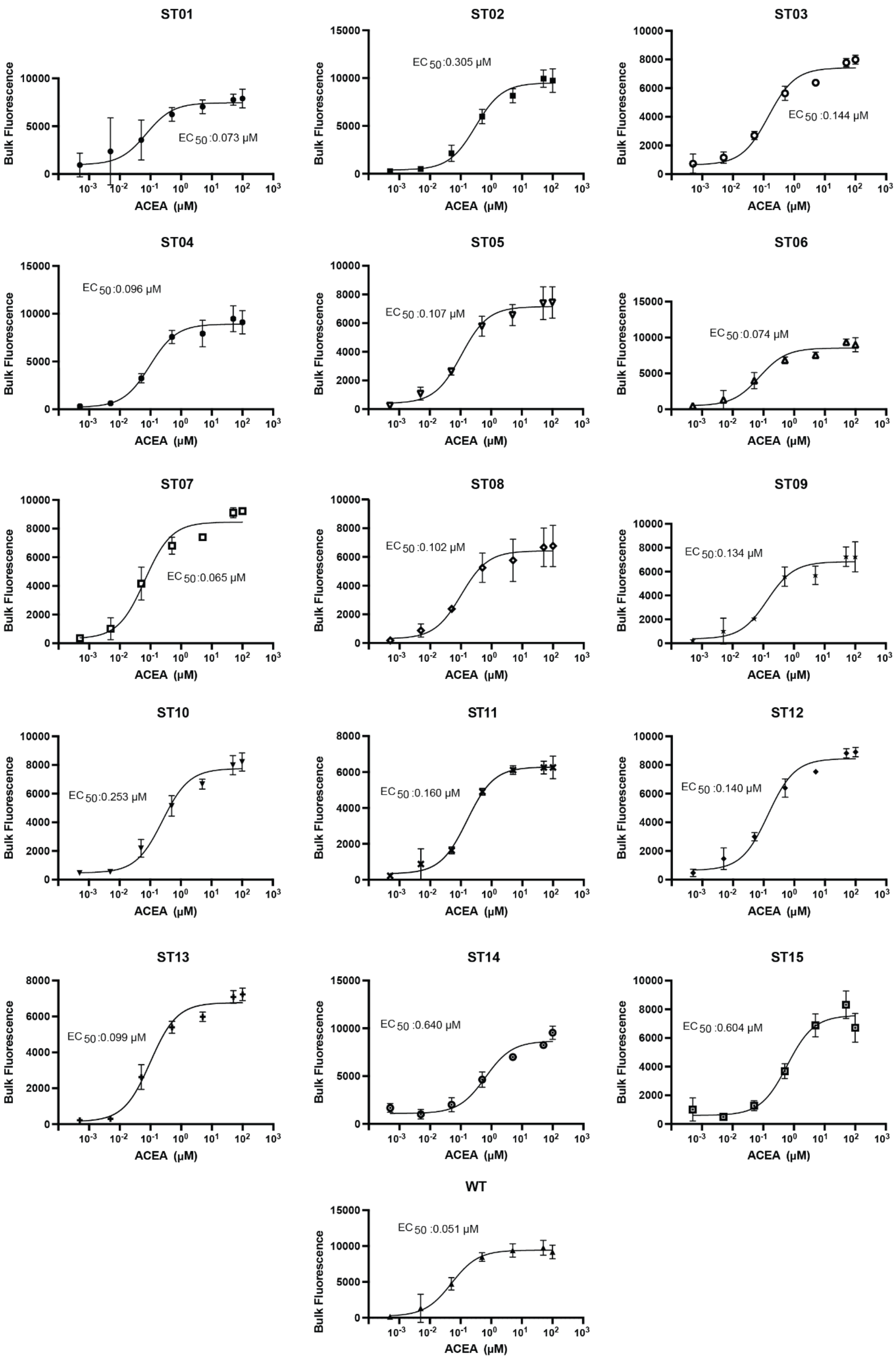


**Supplementary Figure 2. CB1R dose responses with sterol-modified strains.** For all experiments, *n*=3. Nonlinear regression was performed with a 4-parameter logistic equation.

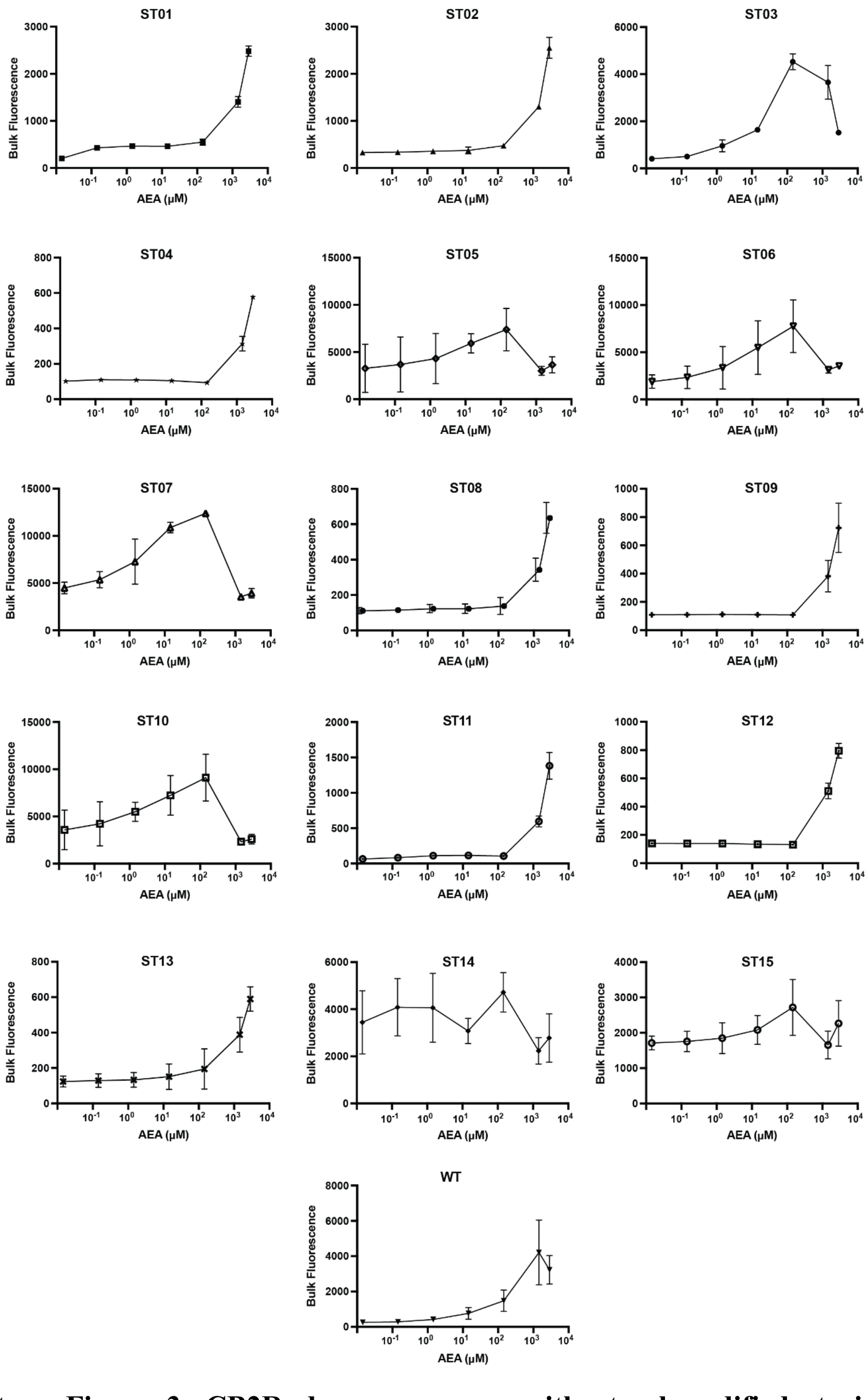


**Supplementary Figure 3. CB2R doses responses with sterol-modified strains.** For all experiments, *n*=3; data points for consecutive doses are connected with lines as a guide to the eye.

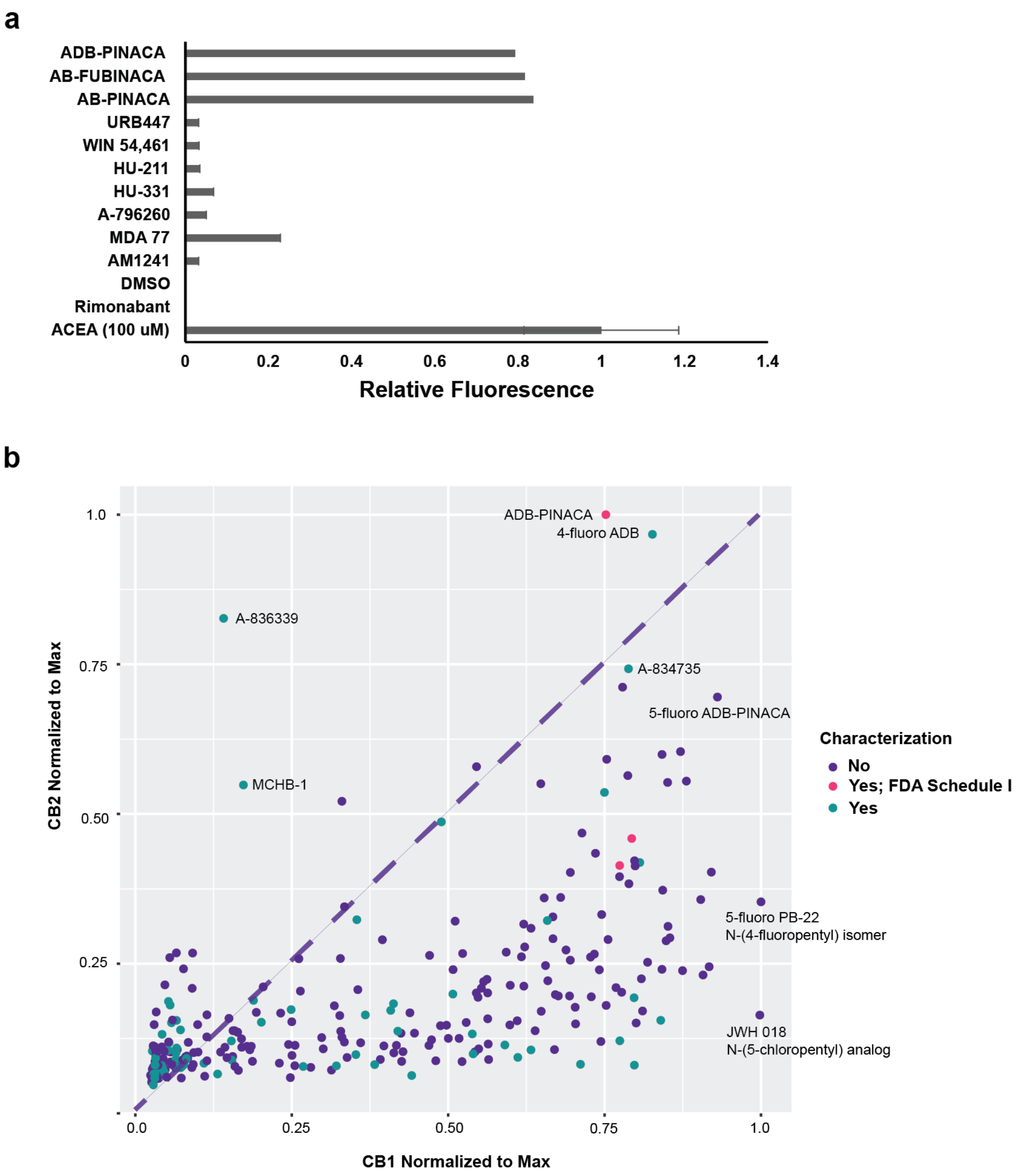


**Supplementary Figure 4. Relative signaling of synthetic cannabinoid compounds at cannabinoid receptors.** (A) Signaling with control compounds at CB1R. Known agonists ADB-PINACA, AB-FUBINACA, and AB-PINACA show strong signaling at 10 nM compared to a positive control (100 µM ACEA). All other compounds (10 nM) are not predicted to activate CB1R. DMSO vehicle was also tested. (B) Synthetic cannabinoid compounds screened against CB1R and CB2R at 10 nM, normalized to the maximum signal for each receptor at this concentration. Large distances from the right and left of the plotted line connote rough bias towards CB1R and CB2R, respectively. Compounds are labeled based on if they, to our knowledge, have been characterized biochemically against one or both cannabinoid receptors.

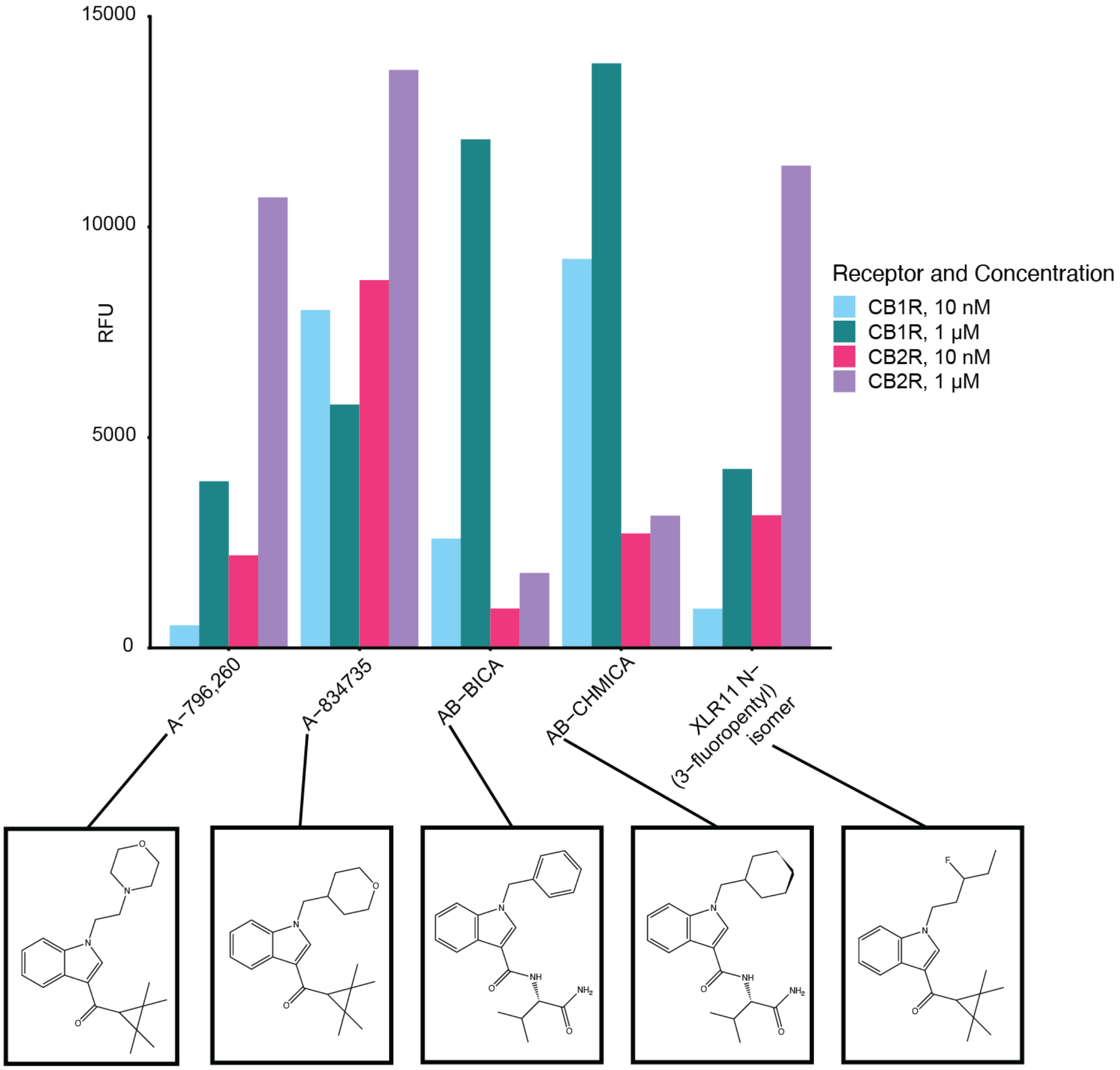


**Supplementary Figure 5. Structure and signaling of isolated synthetic cannabinoid compounds that show biased activity.** (A) Select compounds from the synthetic cannabinoid screen at both CB1R and CB2R. For all experiments, *n*=1.

| Compound | Normalized Activity at 1 μM | Compound | Normalized Activity at 1 μM |
|---|---|---|---|
| AB-PINACA 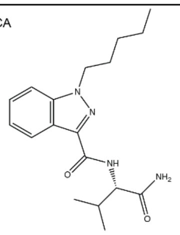 | CB1R: 0.550<br>CB2R: 0.533 | ADB-PINACA 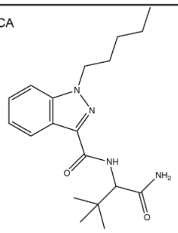 | CB1R: 0.590<br>CB2R: 0.700 |
| AB-PINACA N-(2-fluropentyl) isomer 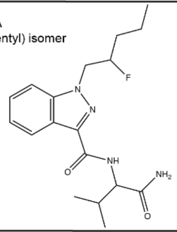 | CB1R: 0.632<br>CB2R: 0.340 | ADB-PINACA isomer 1 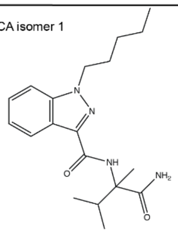 | CB1R: 0.810<br>CB2R: 0.303 |
| AB-PINACA N-(3-fluropentyl) isomer 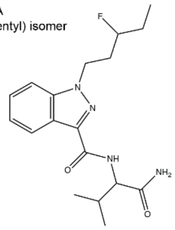 | CB1R: 0.700<br>CB2R: 0.310 | ADB-PINACA isomer 2 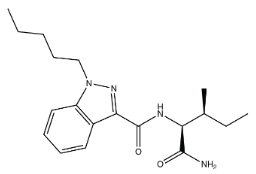 | CB1R: 0.753<br>CB2R: 0.568 |
| AB-PINACA N-(4-fluropentyl) isomer 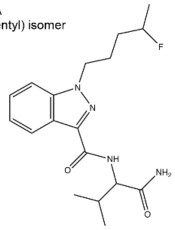 | CB1R: 0.680 | ADB-PINACA isomer 3 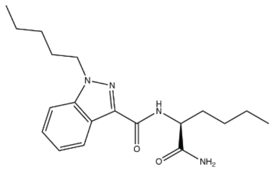 | CB1R: 0.735<br>CB2R: 0.460 |
| ADB-BICA 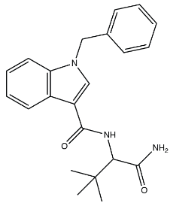 | CB1R: 0.842<br>CB2R: 0.255 | ADB-PINACA isomer 4 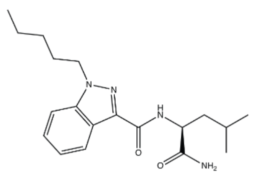 | CB1R: 0.671<br>CB2R: 0.561 |
| ADB-BINACA 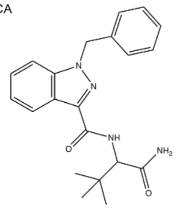 | CB1R: 0.916<br>CB2R: 0.676 | | |

**Supplementary Table 1. Structure and activity of PINACA compounds.**

| 5-fluoro PB-22 isomer | Normalized Activity at 1 μM | 5-fluoro PB-22 isomer | Normalized Activity at 1 μM |
|---|---|---|---|
| 5-fluoro PB-22 3-hydroxyquinoline isomer 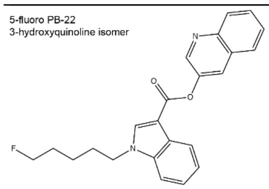 | CB1R: 0.530<br>CB2R: 0.148 | 5-fluoro PB-22 7-hydroxyisoquinoline isomer 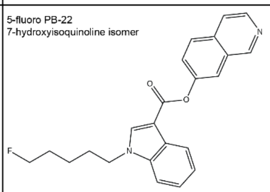 | CB1R: 0.530<br>CB2R: 0.339 |
| 5-fluoro PB-22 4-hydroxyisoquinoline isomer 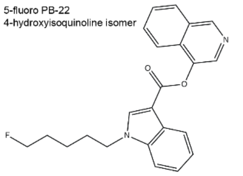 | CB1R: 0.570<br>CB2R: 0.786 | 5-fluoro PB-22 7-hydroxyquinoline isomer 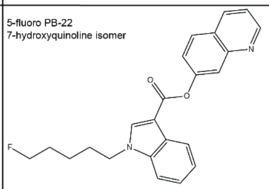 | CB1R: 0.550<br>CB2R: 0.199 |
| 5-fluoro PB-22 4-hydroxyquinoline isomer 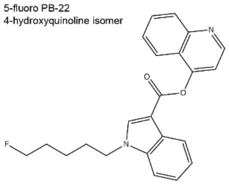 | CB1R: 0.629<br>CB2R: 0.247 | 5-fluoro PB-22 8-hydroxyisoquinoline isomer 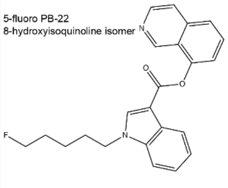 | CB1R: 0.560<br>CB2R: 0.771 |
| 5-fluoro PB-22 5-hydroxyisoquinoline isomer 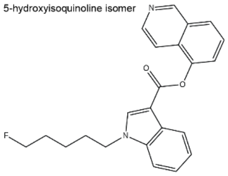 | CB1R: 0.540<br>CB2R: 0.906 | 5-fluoro PB-22 N-(2-fluoropentyl) isomer 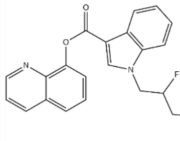 | CB1R: 0.531<br>CB2R: 0.582 |
| 5-fluoro PB-22 5-hydroxyquinoline isomer 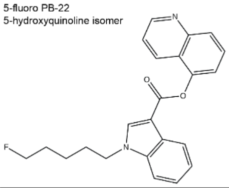 | CB1R: 0.604<br>CB2R: 0.496 | 5-fluoro PB-22 N-(3-fluoropentyl) isomer 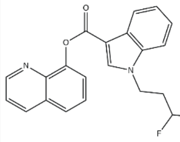 | CB1R: 0.530<br>CB2R: 0.532 |
| 5-fluoro PB-22 6-hydroxyisoquinoline isomer 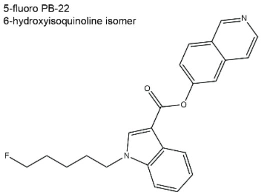 | CB1R: 0.525<br>CB2R: 0.331 | 5-fluoro PB-22 N-(4-fluoropentyl) isomer 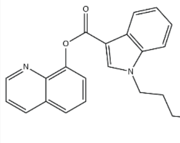 | CB1R: 0.560<br>CB2R: 0.636 |
| 5-fluoro PB-22 6-hydroxyquinoline isomer 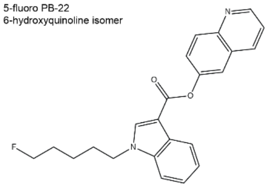 | CB1R: 0.500<br>CB2R: 0.141 | | |

**Supplementary Table 2. Structure and activity of 5-fluoro PB-22 compounds.**
